# Physically informed Monte Carlo simulation of dual-wedge prism-based spectroscopic single-molecule localization microscopy

**DOI:** 10.1101/2023.05.18.541375

**Authors:** Wei-Hong Yeo, Cheng Sun, Hao F. Zhang

## Abstract

**Significance:** The dual-wedge prism (DWP)-based spectroscopic single-molecule localization microscopy (sSMLM) system offers improved localization precision and adjustable spectral or localization performance, but its nonlinear spectral dispersion presents a challenge. A systematic method can help understand the challenges and thereafter optimize the DWP system’s performance by customizing system parameters to maximize spectral or localization performance for various molecular labels.

**Aim:** We developed an MC-based model which predicts the imaging output of the DWP-based sSMLM system given different system parameters.

**Approach:** We assessed our MC model’s localization and spectral precisions by comparing our simulation against theoretical equations and fluorescent microspheres. Furthermore, we simulated the DWP-based system using beamsplitters of Reflectance (R):Transmittance (T) of R50:T50 and R30:T70 and their tradeoffs.

**Results:** Our MC simulation showed average deviations of 2.5 nm and 2.1 nm for localization and spectral precisions against theoretical equations; and 2.3 nm and 1.0 nm against fluorescent microspheres. An R30:T70 beamsplitter improved spectral precision by 8% but worsened localization precision by 35% on average compared to an R50:T50 beamsplitter.

**Conclusions:** The MC model accurately predicted localization precision, spectral precision, spectral peaks, and spectral widths of fluorescent microspheres, as validated by experimental data. Our work enhances the theoretical understanding of DWP-based sSMLM for multiplexed imaging, enabling performance optimization.

## 1 Introduction

Single-molecule localization microscopy (SMLM) allows sub-diffraction-limit imaging of biological structures down to 10 nm^1–3^, with spectroscopic single-molecule localization microscopy (sSMLM) extending this capability to imaging multiple molecular contrasts through analysis of single molecular fluorescence emission spectra^4–10^. In sSMLM, a dispersive component, such as a grating^4–7^ or a prism^8–10^, divides photons from each blinking event to form a spatial image for localization and a spectral image for spectroscopic analysis. Hence, sSMLM can identify each fluorophore based on its characteristic spectrum with nanoscopic spatial precision.

Gratings used in sSMLM are associated with high transmission losses (∼30%), reducing the photon budget, which worsens localization and spectral precisions^11–13^. In contrast, prism-based sSMLM has lower transmission losses but a higher aberration, which causes imaging artifacts^11^. To address these issues, we recently developed a dual-wedge prism (DWP)-based sSMLM that features lower transmission losses, reduced aberrations, the ability to perform highly multiplexed imaging within an extended spectral range, and an adjustable splitting ratio for spatial and spectral imaging^11^. Using DWP-based sSMLM poses challenges, such as a non-uniform spectral dispersion which is poorly understood, but also benefits, such as an adjustable beamsplitter ratio that can allow us to maximize spectroscopic or localization performance.

First, the DWP module provides a higher transmission efficiency than the grating, allowing us to expand our highly multiplexed imaging capabilities through increased localization and spectral precision. Although the DWP module exhibits a nonlinear spectral dispersion, we previously assumed a constant spectral precision in our theoretical analysis of DWP-based sSMLM, since we restricted its operation to a limited spectral range of 650-800 nm^11,12^. When we extend the spectral range to 450-800 nm, which is possible with the DWP due to its higher transmission efficiency throughout the entire spectral range, the spectral dispersion may no longer be considered constant because spectral precision is a function of spectral dispersion^12^.

Next, the splitting ratio of the beamsplitter within the DWP module may be tailored for specific imaging requirements for imaging fluorophores with different emission spectral bandwidths. This is because adjusting the splitting ratio in the DWP module leads to a varying tradeoff between spatial and spectral precisions since the total photon budget is shared between the spatial and spectral channels. While several simulation packages have been developed to help users optimize SMLM imaging conditions and understand SMLM imaging results^14–20^, none of the packages directly simulate the spectra of the fluorophores. Additionally, sSMLM imaging conditions, such as choice of fluorophores and splitting ratios, on the imaging performance quantified by localization and spectral precisions are intrinsically connected but poorly understood in DWP-based sSMLM.

Therefore, there is a need to thoroughly investigate the tradeoff between the extended spectral range and the additional flexibility of variable beamsplitting ratios afforded by the DWP module. In this work, we develop a Monte Carlo (MC) simulation of the 2D-DWP imaging process of individual fluorophore blinking events and determine each fluorophore’s spectral and spatial precisions. Our MC simulation of the 2D-DWP imaging process provides valuable insights into optimizing experimental conditions for imaging, eliminating the need for trial and error. By accurately estimating the localization precision and spectral precision with different splitting ratios, we can directly determine the ideal experimental parameters with different experiments. This optimization enables the use of more fluorophores in a single experiment, unlocking the potential to analyze intricate biological interactions at the nanoscale. The implementation of these refined experimental conditions for multiplexed experiments will enhance our ability to study and understand more complex biological interactions.

## 2 Methods

### 2.1 Image formation process

The MC simulation modeled our current 2D-DWP-based sSMLM system as previously reported^11^. We used two excitation lasers (532 nm, Exlsrone-532-200, Spectra-Physics; and 647 nm, 2RU-VFL-P-2000-647-B1R, MPB Communications) and an inverted microscope body (Ti2-E, Nikon) equipped with a 100× total-internal reflection fluorescence (TIRF) objective (CFI Apochromat TIRF 100XC Oil, Nikon). Figure 1(a) illustrates the photon flow in the detection path. The emitted photons from the fluorescent sample pass through a DWP module, consisting of a beamsplitter prism (BS010 for R50:T50; and BS049 for R30:T70, Thorlabs), a right-angle prism (84-514, Edmund Optics), and a custom-made DWP^11^. The beamsplitter divides the photons into a spatial imaging (0^th^-order) path and a spectral imaging (1^st^-order) path. The right-angle prism then reflects photons in the spatial imaging path, and the custom-made DWP disperses the photons in the spectral imaging path. Finally, an electron-multiplying charge-coupled device (EMCCD) (iXon Ultra 897, Oxford Instruments) simultaneously detects all the photons in the 0^th^-order and 1^st^-order paths for further processing.

**Fig. 1.**
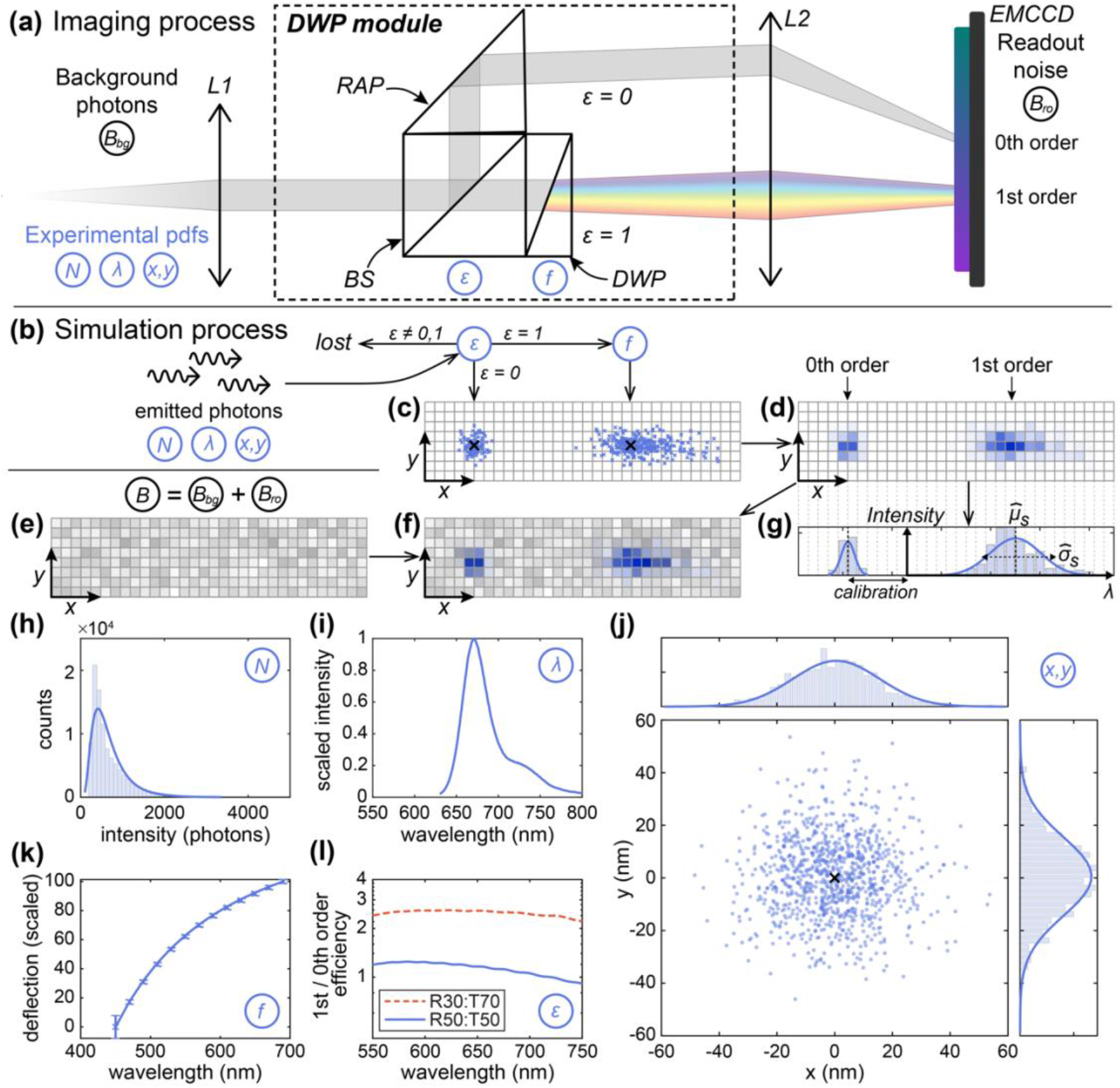
Illustration of the 2D-DWP image formation and MC simulation steps. (a) Schematic of the 2D-DWP image formation. RAP: right-angle prism, BS: beamsplitter. (b) Photon emission simulation process; (c) a simulated image of the photon distribution on the EMCCD following a Gaussian distribution of photon wavelengths; (d) corresponding EMCCD-detected image for photon distribution shown in (c); (e) simulated EMCCD background, including background and readout noises; (f) Combined image from panels d and e; (g) 1D fitting the spectral peak and with from spectral profile, where the 0^th^-order image provides the reference needed to obtain the corresponding 1^st^-order spectral information; (h) measured photon count distribution of AF647 with a 10-ms exposure time; (i) measured bulk emission spectrum of AF647; (j) measured *x* and *y* variation of the individual photons due to the diffraction limit, with *x*- and *y*-summed histograms shown. (k) Measured spectral deflection characteristics of the DWP module. (l) Measured ratios of 1^st^-order / 0^th^-order efficiency in the DWP module when different beamsplitters are used.

### 2.2 MC Simulation process

Figure 1(b) shows our MC simulation process, from generating a blinking event by individual fluorophores to image detection. We start from the ground truth of *N* photons emitted in a single frame from one photoswitching event with parameters *x, y*, and *λ*, where *N* photons are sampled either from a lognormal function or assumed a constant value; *x* and *y* are the spatial positions of each photon; and *λ* is the wavelength of each photon sampled from the emission spectra. Because beamsplitters often have wavelength-dependent splitting ratio and introduce photon losses, we pass the *N* photons through an efficiency function *ε*(*λ*) to determine whether each photon goes to the 0^th^-order path, 1^st^-order path, or is lost. Photons in the 0^th^-order path will be detected directly by the EMCCD; whereas photons in the 1^st^-order path will pass through a dispersion function *f*(*λ*) to determine their wavelength-dependent shift along the *x-*axis (*x*_disp_), mimicking the behavior of the DWP. Next, we combine the locations of the photons, *X*= *x* + *x*_disp_ and Y = y, to determine the exact locations where each photon will be incident on the EMCCD array in Fig. 1(c). This signal is then discretized by individual EMCCD elements due to finite pixel size (a) with varying intensities to give a signal (**S**) illustrated pictorially in Fig. 1(d). We further add a background signal consisting of Poisson-distributed background noises (**B**) and Gaussian-distributed readout noises^21^ (**R**), shown in Fig. 1(e). By summing the digitized signals from Figs. 1(d)-1(e), we applied signal amplification using a Gamma distribution, which is known to effectively model the behavior of an EMCCD^21^. This process yielded *Z*, the simulated sSMLM frame shown in Fig. 1(f).

To obtain the subpixel peak location from the 0^th^-order spatial image, we fit the 0^th^-order image with a bivariate Gaussian function using the maximum likelihood estimation (MLE) method^13^, giving us an estimate of the expected values of the peak locations *x* and *y* as 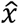 and ŷ, respectively. To obtain the spectral peak from the 1^st^-order spectral image, we first calibrated the *x*-axis of the spectral graph with a spectral calibration procedure (detailed in Sec. 2.4), which allows us to calculate the wavelength values along the *x*-axis for each 0^th^-order localization. We then fitted the spectrally calibrated image with either a univariate or bivariate Gaussian function to estimate the spectral peak 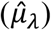and spectral width 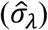 together with the spatial localization. The procedure is illustrated with a 1-dimensional (1D) spectral graph in Fig. 1(g), where we integrated the signal intensity along the *y*-axis for clarity. An alternative method commonly used for spectral characterization is the spectral centroid method, which takes the weighted average of the different wavelengths detected to calculate a spectral centroid associated to each localization^11,12,22^. In this study, we fitted the resulting image with a bivariate Gaussian function. This approach enhances noise rejection by taking advantage of the larger number of samples available in 2D data, as opposed to summing it along the *y*-axis in 1D. As the 2D Gaussian fit utilizes information from both dimensions, it provides a more accurate representation of the underlying distribution and consequently improves the robustness against noise.

To illustrate the functions *N, λ, f*(*λ*) and *ε*(*λ*) in Table 1 used in the simulation, we extracted the distribution of photon count and spectral signature of the commonly used SMLM dye Alexa Fluor 647 (AF647). Fig. 1(h) shows the lognormal fitted distribution of photon counts *N* of AF647 extracted using the process described in Sec. 2.3.1. Fig. 1(i) shows the spectral distribution *λ* of AF647 obtained with the procedure described in Sec. 2.3.2, and Fig. 1(j) shows a simulated point spread function (PSF) using a 2D-Gaussian function, which is a close approximation to the Airy disc^23^. Fig. 1(k) shows the spectral dispersion function, *f*(*λ*), experimentally measured using the procedure described in Sec. 2.3.3. Fig. 1(l) shows the efficiency function, *ε*(*λ*), for beamsplitters with reflectance (R):transmittance (T) of R30:T70 and R50:T50 experimentally measured using the procedure described in Sec. 2.3.4. Finally, we repeated the process 20,000 times to obtain the spatial and spectral precisions, defined as the standard deviation of 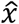 and 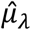 respectively. We summarized the parameters used in the simulation in Table 1.

**Table 1.** Summary of simulation parameters.

| Parameter | Meaning | Value/Definition |
| --- | --- | --- |
| *Vectors describing each photon in a single localization event* |  |  |
| $x, y$ | vector of calculated $x$ and $y$ positions | - |
| $\lambda$ | vector of wavelengths | - |
| $\theta$ | vector of the paths taken by the photons after the beamsplitter | - |
| $x_{\text{disp}}$ | vector of $x$ dispersions due to the DWP module | - |
| *Matrices describing each pixel in the EMCCD pixel array* |  |  |
| **S** | the signal on the pixel array | **S** ; $S_{i,j} = \sum_{i,j}(x, y)$ |
| **B** | background noise on the pixel array | **B** $\sim \text{Po}(b_{\text{bg}})$ |
| **R** | readout noise on the pixel array | **R** $\sim \text{N}(0, b_{\text{ro}}) \times A / \text{ADU}$ |
| **Z** | output on the EMCCD pixel array | **Z** $= \Gamma(\mathbf{S} + \mathbf{B}, A) / \text{ADU} + \mathbf{R}$ |
| **Z'** | output on the EMCCD pixel array after background subtraction | **Z'** $= \mathbf{Z} - A / \text{ADU} \times b_{\text{bg}}$ |
| *Functions used in the simulation* |  |  |
| $N$ | p.d.f. of the number of photons from fluorophore | $N \sim \text{Lognormal}(\mu, \sigma^2)$ |
| $\lambda$ | p.d.f. of the wavelength of photon from fluorophore spectrum | Custom function |
| $\varepsilon(\lambda)$ | efficiency function of DWP module | $\varepsilon(\lambda) \in \{0, 1, -1\}$ |
| $f(\lambda)$ | spectral dispersion function of DWP module | $f(\lambda) = x_{\text{disp}}$ |
| *Constants used in this paper* |  |  |
| $N$ | number of photons from fluorophore | 2000 |
| $b_{\text{bg}}$ | mean number of background photons per pixel | 15 |
| $b_{\text{ro}}$ | standard deviation of readout noise | $1.9 \text{ e}^-$ |
| $A$ | gain on the EMCCD output | 100 |
| ADU | analog to digital unit on the EMCCD | 13.6 |
| $a_x, a_y$ | pixel size | $16 \mu\text{m}/\text{px}$ |
| $a_\lambda$ | spectral dispersion (linear equivalent) | $4.3 \text{ nm}/\text{px}$ |
| NA | numerical aperture | 1.49 |
| $F_{\text{EM}}$ | electron multiplying factor | $\sqrt{2}$ |

### 2.3 Extraction of parameters from experimental data

#### 2.3.1 Distribution of photon counts

To obtain physical values for the simulation, we extracted key parameters used in our MC simulation from our sSMLM experimental data. Fig. 1(h) shows the histogram of the photon count *N* of AF647 molecules imaged at an exposure time of 10 ms. The photon counts were determined using ThunderSTORM^24^, and the values obtained were fitted with a lognormal distribution^25^.

#### 2.3.2 Spectral distribution of fluorophores/microspheres

The example spectral signature of AF647 shown in Fig. 1(i) can usually be obtained from open-source databases, such as FPbase^26^. For data unavailable in FPbase, such as for microspheres tested in this work, we perform a Vis-IR measurement of the emission spectra using the Nanodrop 3300 fluorospectrometer, which provides the bulk emission spectra of the microspheres.

#### 2.3.3 Spectral dispersion of the DWP module

We experimentally determined the spectral dispersion of the DWP module using a supercontinuum laser with an acousto-optic tunable filter (AOTF) to precisely adjust the output wavelength. We measured the angular deflection of the beam of the DWP module compared to the measurements without the DWP module inserted at different wavelengths. The experimental angular deflection curve was validated by the theoretical curve given by the Sellmeier equation^27^, as shown in Fig. S1(a), with a good agreement. To better fit the theoretical curve, we used a rational function and found that the following equation works satisfactorily from 400 nm to 900 nm:

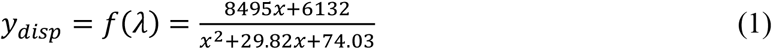

with 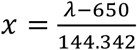as a scaling factor. Since Eq. (1) is an invertible rational function, we can find the inverse equation:

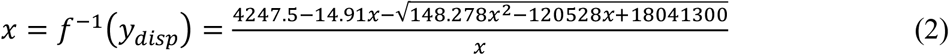

with λ = 144.342*x* + 650 to obtain the wavelength from the angular deflection. Eq. (2) adequately describes the scaling and translation optical transformations associated with using the DWP module. This is because, after any beam passes through the DWP module, the only transformations it can undergo are scaling (when the pathlength between the DWP module and the camera plane is changed) or translation (when the pathlength between the 0^th^- and 1^st^-order on the camera plane is changed). Following this, a linear equation between the linear dispersion and the photon wavelength can describe any spectral calibration curve of the DWP module

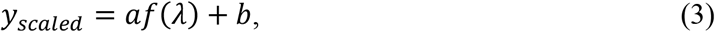

where *a* is the scaling factor and *b* is the translation factor. We use Eq. (3) to compute the spectral window of interest in the pixel for each localization detected in the spatial image. However, when we need to compute the spectral peak or spectral width, we need to combine the results of Eq. (2) and the inverse of Eq. (3) to get:

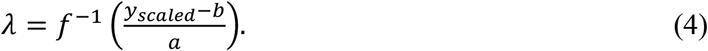

We compared the theoretical and the empirical curve in Fig. S1(b) in the supplemental notes and found the angular error between the theoretical and the empirical curve to be less than 0.3%.

#### 2.3.4 Optical transmission efficiency of the DWP module

We evaluated the optical transmission efficiencies of the beamsplitter, right-angle prism, and DWP by measuring the transmitted and reflected optical power ratios using a supercontinuum laser with an AOTF setup, as described in Sec. 2.3.3. We then calculated the efficiencies at different wavelengths by calculating the ratios of transmitted (*P*_*T*_) and reflected (*P*_*R*_) optical power to the total incident power (*P*_*I*_). Since our AOTF adjustable spectral range was from 450 nm to 690 nm, we used the data to validate the Thorlabs datasheet for the beamsplitters. To extend the spectral range beyond 690 nm for the right-angle prism, we assumed that the efficiencies remain relatively constant. We utilized the average efficiency value measured for the range of 450 nm to 690 nm and applied it throughout our simulated spectral range of 450 nm to 800 nm. For the DWP, we extrapolated our results for efficiencies beyond 690 nm. In Fig. S2, we present the experimentally determined plots as dots for each optical component, and the corresponding dotted lines represent the simulated curves.

### 2.4 Post-processing of EMCCD images

#### 2.4.1 Spectral calibration

To calibrate the spectral dispersion of the DWP, we used a custom-made nanohole array as a static target for spectral calibration and five bandpass filters (Table S1). The nanohole array (Fig. S3) was fabricated on a gold-plated glass coverslip using a Helios Nanolab 600 Focused ion beam-scanning electron microscope. To obtain the average spectral dispersion (*x*_disp_) of each hole at different wavelengths, we imaged the nanohole array through the five bandpass filters which have center wavelengths of 532 nm, 605 nm, 635 nm, 685 nm, and 750 nm. Based on these measurements, we performed a linear fit using Eq. 4 to determine the spectral deflection at different wavelengths of the DWP module.

#### 2.4.2 Correcting for nonlinearity in spectral fits in the DWP module

To accurately fit the spectra obtained from the DWP module, it is crucial to consider the inherent nonlinearity in spectral dispersion. As shown in Fig. 2(a), a hypothetical light source with uniform spectral power density undergoes spectral dispersion after passing through the DWP module, resulting in non-uniform intensity across the image sensor, as shown in Fig. 2(b). This effect is particularly noticeable in regions with lower spectral dispersion, where the incident power is greater. Consequently, fitting a Gaussian curve to the spectrum would result in a bias towards the longer wavelengths due to the higher intensity in the region. To account for this spectral bias, we normalized the spectra with the spectral dispersion, as illustrated in Fig. 2(c), which results in a corrected image with uniform intensity throughout the image sensor. This normalization process corrected the nonlinear spectral dispersion in the DWP module and ensured accurate spectral fitting.

**Fig. 2.**
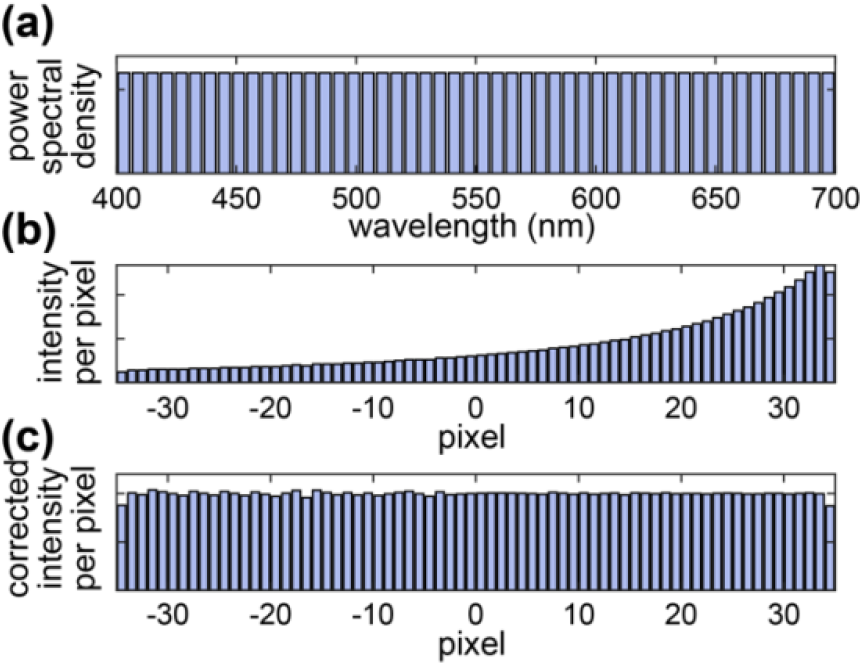
Illustration of the correction process of nonlinearity in spectral dispersion with DWP. (a) Illustration of the wavelength distribution of an ideal uniformly distributed spectral density light source; (b) Power density detected by the EMCCD after the DWP; (c) corrected power density.

### 2.5 Validation with fluorescence microspheres

To validate our MC model, we imaged microspheres with emission peaks at 560 nm, 580 nm, 605 nm, 645 nm, and 720 nm (F8800, F8794, F8801, F8806, and T8870, Invitrogen) deposited on #1.5-thick glass coverslips (12-541-B, Fisher Scientific). The other physical properties of these microspheres are detailed in Table S2. We first washed the glass coverslips with PBS and coated the coverslips with 0.001% poly-L-lysine solution for 5 min. We then washed the coverslips with PBS and stained the glass coverslips with a diluted sample of each microsphere for 1 hour. Next, we washed off the unbounded microspheres with PBS three times and added a single drop of Antifade buffer (P36961, Invitrogen). Finally, we sealed the coverslips with black nail polish (114-8, Ted Pella) and left them overnight to dry.

We first imaged the microspheres without the DWP module to obtain the photon count of each nanosphere, then imaged the same microspheres with the DWP module for 100 frames. To obtain the spatial localization precision, we processed the 100 frames of microspheres using ThunderSTORM^24^ and treated the standard deviation from 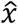 calculated from the 100 frames as the localization precision. To obtain the spectral precision, we identified a region of interest from 450 nm to 800 nm relative to the 0^th^-order image after spectral calibration using the method previously described^5,11,22^. This region of interest gives us a spectral profile similar to the one seen in Fig. 1(f), which we can use to analyze the spectral signature of the fluorophore further. Next, we subtracted the background using averaged neighboring windows of the 1^st^-order image. Then, we performed a least-squares fit of the spectral profile with a bivariate Gaussian function to identify the spectrum corresponding to the 0^th^-order blinking.

## 3 Results and Discussion

### 3.1 Validating MC simulation with theoretical analyses

To validate our MC model, we first benchmarked the precisions generated from our model against theoretical models of spatial localization precision Δ*x* defined as^13^:

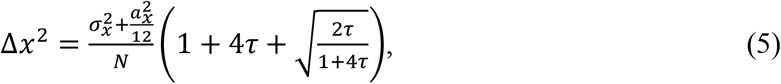

where the background correction factor τ is defined as 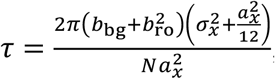 and spectral precision Δμ_λ_ is defined as^12^:

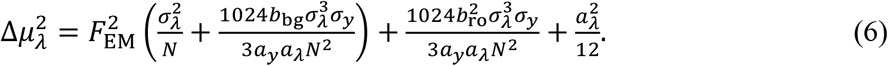

Eq. 5 describes the performance limit of the MLE when fitting the PSF into an approximate 2D Gaussian curve, also referred to as the Cramér-Rao lower bound^13^. Eq. 6 assumes (1) a uniform spectral dispersion, (2) a Gaussian spectral shape with a spectral peak of μ_λ_ and a spectral width of σ_λ_, and (3) spectral centroid in the calculation of spectral precision^12^. The spectral centroid method takes the weighted average of the intensities captured in the pixels associated with different wavelengths, yielding a weighted representation of the spectra recorded.

To provide a fair comparison to validate our simulation, we matched the simulation based on the assumptions of Eq. 6 listed above. We computed the spatial localization precision, defined as the standard deviation of the localized spot from the true location, and the spectral precision, defined as the standard deviation of the spectral centroid or the fitted spectral peak. We performed MC simulations under the following conditions: *N* = 700, *b*_bg_ = 15, spectral peak = 680 nm, and spectral full-width-at-half-maximum (FWHM) = 41 nm, using a linear spectral dispersion of 5 nm/px, unless otherwise stated.

Figure 3 shows the influence of photon counts *N*, background photons *b*_bg_, and emission spectral peak μ_λ_ on lateral localization and spectral precisions. In Figs. 3(a)-3(c), we compared the theoretical localization precisions calculated with Eq. 5 to our MC simulation, with mean absolute errors of 2.5 nm, 0.8 nm, and 0.3 nm relative to the theoretical values, respectively. Similarly, in Figs. 3(d)-3(g), we compared the theoretical spectral precisions calculated with Eq. 6 to our MC simulation using the spectral centroid method, with mean absolute errors of 2.1 nm, 7.6 nm, and 2.2 nm, respectively. For the spectral fitting method, the simulated precisions are consistently better, with mean absolute errors of 6.3 nm, 4.6 nm, and 3.6 nm for Figs. 3(d-g), respectively.

**Fig. 3.**
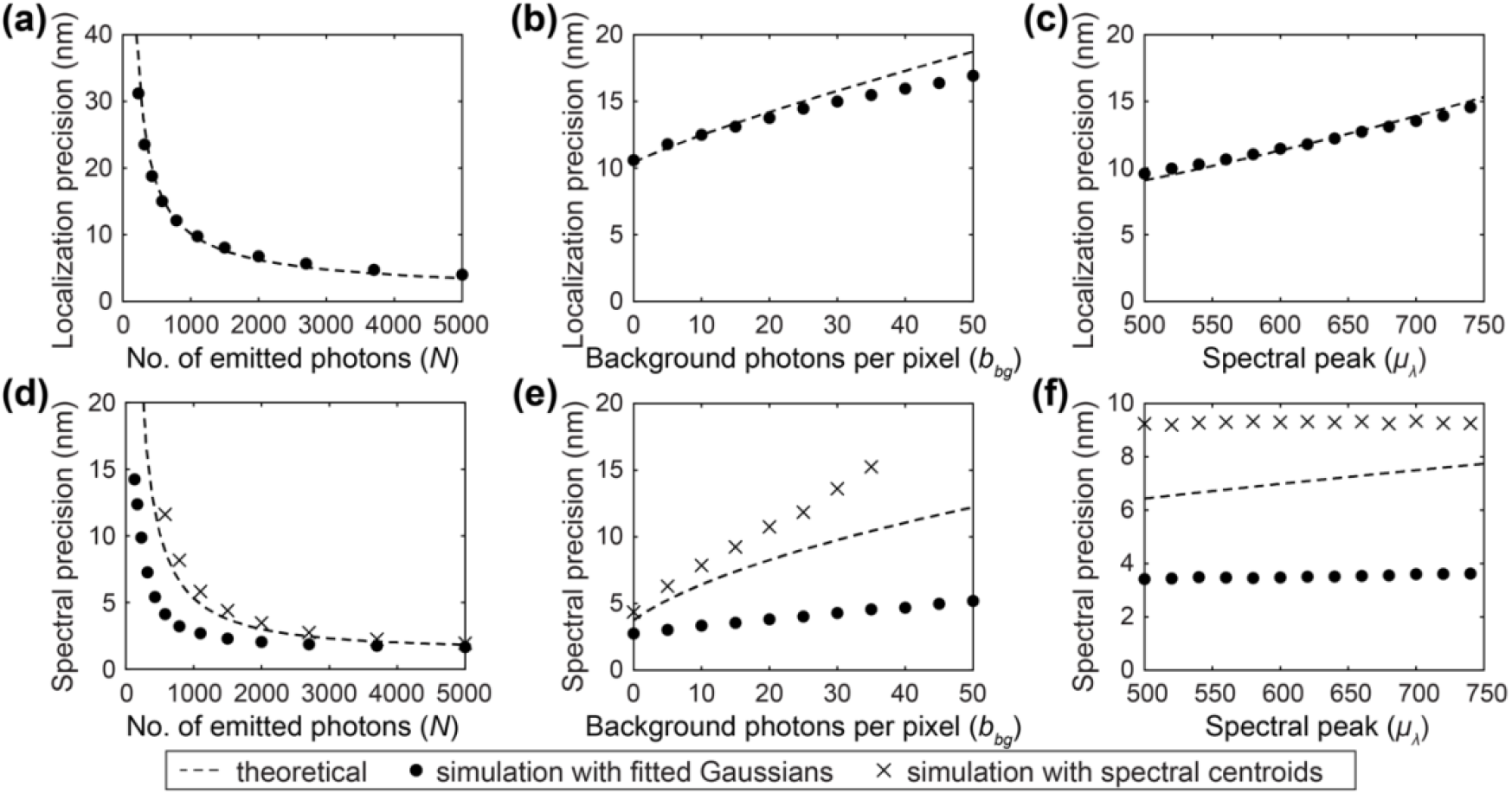
Comparison of localization precisions between theoretically predicted results (dashed line) and simulation results (dots) with respect to (a) photon count, (b) background photons, and (c) emission spectral peak. Comparison of spectral precisions between theoretically predicted results (dashed lines), simulated results calculated with spectral centroids (crosses), and simulated results calculated with a bivariate Gaussian fit (dots) with respect to (d) photon count, (e) background photons, and (f) spectral peak.

In Figs. 3(a) and 3(d), we observe that increasing the number of emitted photons from 1500 to 5000 improved the localization precision from 8.1 nm to 4.0 nm. Similarly, the spectral precision improved from 4.4 nm to 2.0 nm with the spectral centroid method and 3.2 nm to 1.6 nm with the spectral fitting method. With a limited photon budget, splitting the photon budget between 0^th^-order and 1^st^-order images can affect the tradeoff between localization precision and spectral precision.

In Figs. 3(b) and 3(e), increasing background photons from 0 to 30 per pixel worsens localization precision from 10.6 nm to 15.0 nm. Likewise, spectral precision worsens from 4.4 nm to 13.6 nm with the spectral centroid method and from 2.7 nm to 4.3 nm with spectral fitting. The spectral fitting method is more robust against noise because it estimates the background term by fitting for and averaging over all the pixels available in the spectral window, reducing the variation in the estimated spectral peak 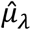. In contrast, the spectral centroid method takes the weighted average of the spectral window, which is more susceptible to noise variations. Additionally, accurately estimating the background level can be challenging^28^, which can result in biases to the spectral centroid value. A positive bias can result from the presence of background contribution to the spectral window, which can shift the spectral centroid value towards the mean of the spectral window and reduce the spectral precision. Hence, we prefer the bivariate Gaussian fitting method due to its robustness in estimating the spectral peak 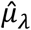 and calculating for spectral precisions.

In Figs. 3(c) and 3(f), we observe that as the center wavelength of the emission peak increases from 500 nm to 740 nm, the localization precisions deteriorate from 9.6 nm to 15.0 nm. Simultaneously, the spectral precision is stable at approximately 9.2 nm with the spectral centroid method and at 3.5 nm with the spectral fitting method. This effect is more pronounced in Fig. 3(c), where the wider FWHM of the PSF results in improved localization precision. However, in Fig. 3(f), a spectral emission FWHM of 47 nm plays a more significant role in determining the overall width of the 1^st^-order image with added spectral dispersion. Overall, our MC model agrees well with the theoretical predictions, with a worse-case average absolute error of 2.5 nm for lateral precision and 2.1 nm for spectral precision with the spectral centroid method across all cases.

### 3.2 MC simulation of different beamsplitters

After validation, we used the MC model to predict the performance of different beamsplitters used in the DWP module. Fig. 4 shows how photon counts *N*, background photons *b*_bg_, and emission spectral peak μ_λ_, affect spatial localization and spectral precisions with different beamsplitters. We conducted MC simulations under the following conditions: *N* = 700, *b*_bg_ = 15, μ_λ_ = 680 nm, emission spectral FWHM = 41 nm, and we used the DWP system with different beamsplitter ratios unless otherwise stated.

**Fig. 4.**
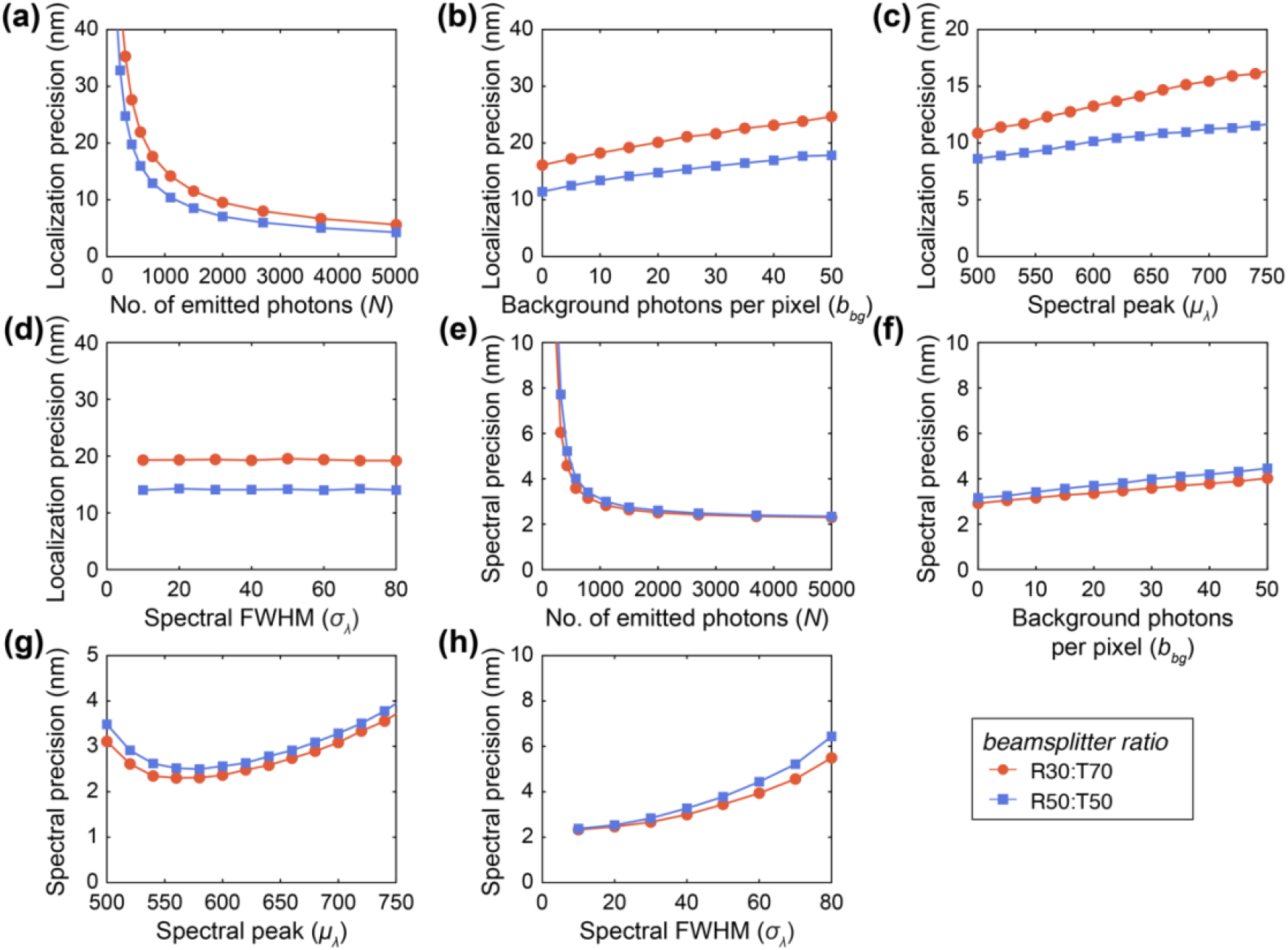
Comparison of localization precisions between an R50:T50 (orange dots) and R30:T70 (blue squares) beamsplitter with respect to (a) photons count, (b) background photons, (c) spectral peak, and (d) spectral FWHM. Comparison of spectral precisions between an R50:T50 (orange dots) and R30:T70 (blue squares) beamsplitter with respect to (e) photons count, (f) background photons, (g) spectral peak, and (h) spectral FWHM.

Figures 4(a)-4(d) show that using an R30:T70 beamsplitter results in a 35% worse localization precision on average compared to an R50:T50 beamsplitter. However, Figs. 4(e)-4(h) demonstrate that an R30:T70 beamsplitter provides 8% better spectral precision on average compared to an R50:T50 beamsplitter. This is because the R30:T70 beamsplitter redirects more of the photon budget towards the spectral channel, which leads to lower spectral precisions, as shown in Fig. 3(d). The tradeoff between spectral and spatial localization precisions is especially prominent in photon-limited photoswitching events when *N* < 1000, as shown in Figs. 4(a) and 4(e), where we observed up to a 12.3 nm worsening in localization precision and up to 2.7 nm improvement in spectral precision for *N* = 320. From Fig. 1(h), many photoswitching events of AF647 have *N* < 1000, which suggests that the change of beamsplitter can play a significant role in optimizing the accuracy in the classification of different fluorophores.

Increasing the number of background photons from 0 to 50 per pixel leads to an increase in the localization and a minimal increase in the spectral precision in both beamsplitters. In Fig. 4(b), the localization precision worsens from 11.4 nm to 17.8 nm for the R50:T50 beamsplitter and from 16.1 nm to 24.7 nm for the R30:T70 beamsplitter. In Fig. 4(f), the spectral precision worsens slightly from 3.2 nm to 4.5 nm for the R50:T50 beamsplitter and from 2.9 nm to 4.0 nm for the R30:T70 beamsplitter. Overall, switching from an R50:T50 beamsplitter to an R30:T70 beamsplitter led to an average worsening in localization precision by 5.6 nm and an improvement in spectral precision by 0.3 nm when the number of background photons per pixel is varied.

Figure 4(c) shows that as the spectral peak changes from 500 nm to 750 nm, the localization precision worsens from 8.6 nm to 11.8 nm for the R50:T50 beamsplitter, and from 10.9 nm to 16.6 nm for the R30:T70 beamsplitter. This worsening is attributed to the change in PSF’s FWHM, as observed in Fig. 3(c). However, Fig. 4(g) reveals a nonlinear relationship between the spectral precision and spectral peak, unlike in Fig. 3(f), with mean values of 3.0 nm for the R50:T50 beamsplitter and 2.8 nm for the R30:T70 beamsplitter. This nonlinearity results from the nonlinear spectral dispersion of the DWP system, as shown in Fig. 1(k), causing spectral precision to vary based on the emission wavelength. Overall, switching from an R50:T50 beamsplitter to an R30:T70 beamsplitter led to an average worsening in localization precision of 3.5 nm, an average improvement in spectral precision of 0.2 nm, and up to a 0.4 nm improvement in spectral precision at a spectral peak of ∼540 nm.

Figure 4(d) shows that the localization precision remains relatively stable when the FWHM spectral bandwidth increases from 10 nm to 80 nm. Figure 4(h) shows that the spectral precision worsens from 2.3 nm to 6.4 nm for the R50:T50 beamsplitter and from 2.3 nm to 5.5 nm for the R30:T70 beamsplitter. This increase is due to the broader spread of the 1^st^-order image on the EMCCD, with results in a lower signal-to-noise ratio in the detected spectra for a fixed photon count *N*, and, consequently, poorer precision. However, localization precision remains mostly unaffected, as the PSF does not change significantly compared to the spectral shape. Overall, switching from an R50:T50 beamsplitter to an R30:T70 beamsplitter led to an average worsening in localization precision by 5.2 nm and an average improvement of spectral precision by 0.4 nm.

### 3.3 Testing MC simulation against experimental data

Figure 5 tests our MC simulation against experimental results obtained by imaging microspheres. The results demonstrate a good match between the MC-DWP simulation and our experimental data for localization precision, as shown in Figs. 5(a)-5(r), with an average absolute error of 2.7 nm for the R30:T70 beamsplitter and 1.5 nm for the R50:T50 beamsplitter. However, for spectral precision shown in Figs. 5(c)-5(t), we observed a slightly higher error, with an average of 0.9 nm for the R30:T70 beamsplitter, and 1.0 nm for the R50:T50 beamsplitter, compared to the simulation.

**Fig. 5.**
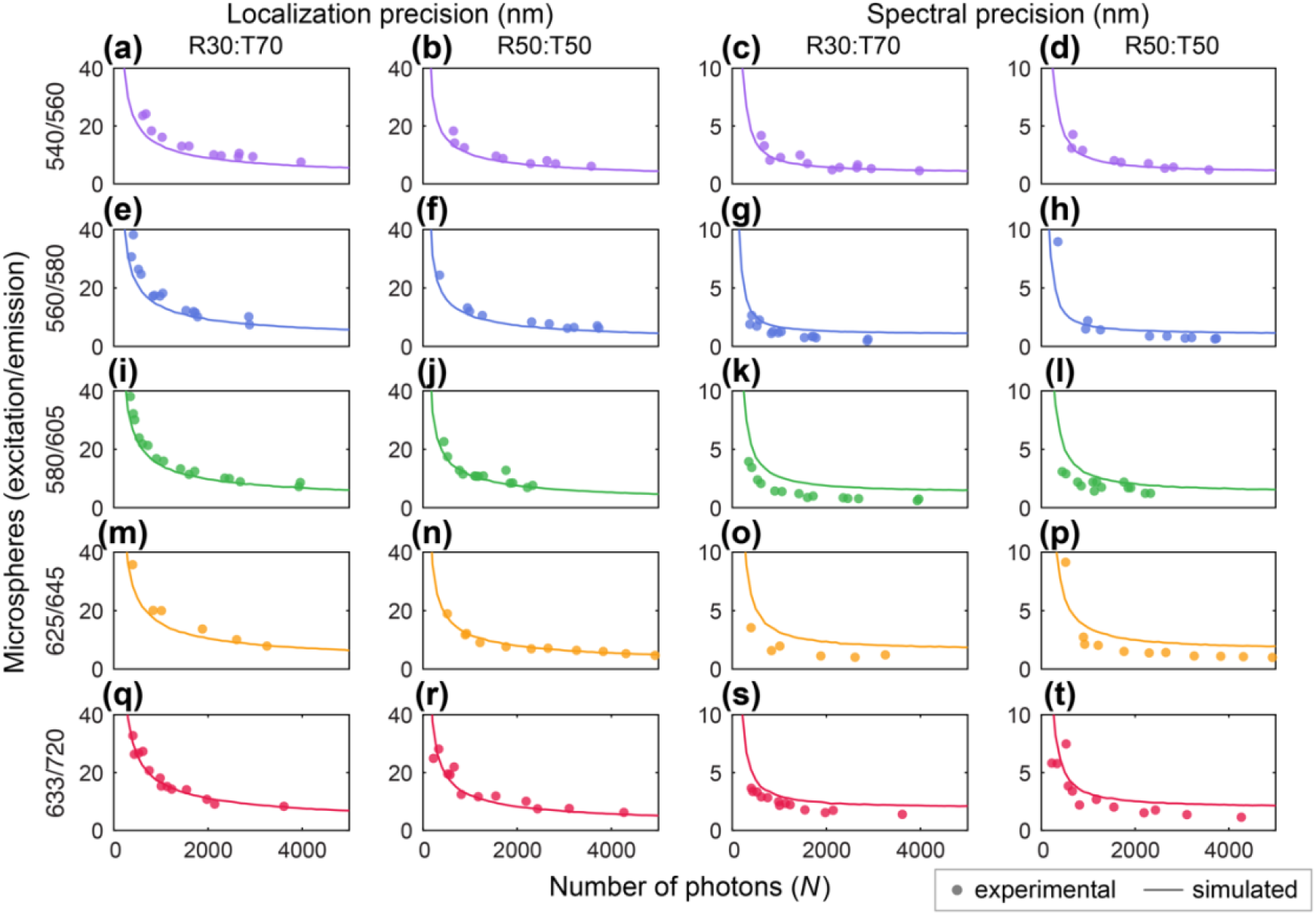
Localization and spectral precisions of different microspheres under varying numbers of emitted photons with DWP-based sSMLM system. Each row shows the localization and spectral precisions of a single species of microspheres. The two columns to the left (a)-(r) are localization precisions, whereas the two columns to the right (c)-(t) are spectral precisions. The beamsplitter used in each case is indicated at the top of each column.

The difference in spectral precision can be attributed to two factors: (1) spectral variations within individual microspheres and (2) imperfect background estimation in the experimental results. While the simulation results used the bulk spectrum of the microspheres to generate the data, the experimental results measured the spectra of individual microspheres. At the molecular level, individual spectra are not readily available, and it is known that when adding two unequal Gaussian probability density functions (PDFs), the result is a PDF with a broader width. Because the individual spectral signatures of the microspheres differ from the bulk phase due to heterogeneity^22^, their narrower widths likely collectively combine to produce a broader bulk phase spectral signature. As shown in Fig. 4(h), increasing the spectral FWHM by 10 nm results in an approximate 10% degradation in spectral precision. The influence of spectral heterogeneity present in our experimental findings explains the slightly improved calculated spectral precision in the experiments compared to the simulation results.

Furthermore, background estimation and subtraction can be challenging and biased because the microspheres are not photoswitching. During data processing, background subtraction may have caused some microspheres to have consistently lower spectral precisions than the simulated values, resulting in systematic errors in the calculated spectral precision^28^. Nonetheless, our MC-DWP simulation provides a reasonable estimate of the performance of the different nanospheres.

## 4 Conclusion

We developed a physically informed MC model for predicting the imaging performance of DWP-based sSMLM systems. Our model accurately predicts both localization and spectral precisions, as well as spectral peaks and widths in fluorescent microspheres, which provides a theoretical foundation for optimizing the performance of multiplexed imaging. By simulating the DWP-based system using different beamsplitters, we found that an R30:T70 beamsplitter can reduce the spectral precision by up to 14%, albeit with a penalty of 35% in localization precision on average. Our work can guide the optimization of imaging parameters for common fluorophore combinations to maximize both localization and spectral performance. Moreover, our model can generate ground-truth data for different fluorophores, which could improve machine-learning algorithms for more accurate fluorophore identification^5,29^. While there are limitations in the agreement between our model and experimental data due to the spectral heterogeneity of individual microspheres, we are exploring new theoretical and simulation models that can potentially describe the spectral heterogeneity of microspheres and fluorophores, which could further improve imaging performance and accuracy.

## Acknowledgments

The authors sincerely acknowledge the generous support from NIH grants R01GM139151, R01GM140478, U54CA268084, and R01GM143397, and NSF grants CHE-1954430 and EFMA-1830969. This work used the EPIC facility of Northwestern University’s NU*ANCE* Center, which has received support from the SHyNE Resource (NSF ECCS-2025633), the IIN, and Northwestern’s MRSEC Program (NSF DMR-1720139). Wei-Hong Yeo is supported by the Christine Enroth-Cugell Fellowship for Vision and Neuroscience at Northwestern University.

## Code, Data, and Materials Availability

The source code for the Monte Carlo simulation can be found in the following link in GitHub:https://github.com/FOIL-NU/MC-DWP

## References

1. E. Betzig et al., “Imaging Intracellular Fluorescent Proteins at Nanometer Resolution,” Science 313(5793), 1642–1645 (2006) [doi:10.1126/science.1127344].

2. M. J. Rust, M. Bates, and X. Zhuang, “Sub-diffraction-limit imaging by stochastic optical reconstruction microscopy (STORM),” Nat Methods 3(10), 793–796 (2006) [doi:10.1038/nmeth929].

3. R. Jungmann et al., “Multiplexed 3D cellular super-resolution imaging with DNA-PAINT and Exchange-PAINT,” Nat Methods 11(3), 313–318 (2014) [doi:10.1038/nmeth.2835].

4. B. Dong et al., “Super-resolution spectroscopic microscopy via photon localization,” Nat Commun 7(1), 12290 (2016) [doi:10.1038/ncomms12290].

5. Z. Zhang et al., “Machine-learning based spectral classification for spectroscopic single-molecule localization microscopy,” Opt. Lett., OL 44(23), 5864–5867, Optica Publishing Group (2019) [doi:10.1364/OL.44.005864].

6. K.-H. Song et al., “Three-dimensional biplane spectroscopic single-molecule localization microscopy,” Optica 6(6), 709 (2019) [doi:10.1364/OPTICA.6.000709].

7. K.-H. Song et al., “Symmetrically dispersed spectroscopic single-molecule localization microscopy,” Light Sci Appl 9(1), 92 (2020) [doi:10.1038/s41377-020-0333-9].

8. Y. Suzuki et al., “Imaging of the fluorescence spectrum of a single fluorescent molecule by prism-based spectroscopy,” FEBS Letters, 5 (2002).

9. Z. Zhang et al., “Ultrahigh-throughput single-molecule spectroscopy and spectrally resolved super-resolution microscopy,” Nat Methods 12(10), 935–938 (2015) [doi:10.1038/nmeth.3528].

10. M. J. Mlodzianoski et al., “Super-Resolution Imaging of Molecular Emission Spectra and Single Molecule Spectral Fluctuations,” PLoS ONE 11(3), M. Sauer, Ed., e0147506 (2016) [doi:10.1371/journal.pone.0147506].

11. K.-H. Song et al., “Monolithic dual-wedge prism-based spectroscopic single-molecule localization microscopy,” Nanophotonics 11(8), 1527–1535, De Gruyter (2022) [doi:10.1515/nanoph-2021-0541].

12. K.-H. Song et al., “Theoretical analysis of spectral precision in spectroscopic single-molecule localization microscopy,” Review of Scientific Instruments 89(12), 123703 (2018) [doi:10.1063/1.5054144].

13. B. Rieger and S. Stallinga, “The Lateral and Axial Localization Precision in Super-Resolution Light Microscopy,” ChemPhysChem 15(4), 664–670 (2014) [doi:10.1002/cphc.201300711].

14. D. Sage et al., “Super-resolution fight club: assessment of 2D and 3D single-molecule localization microscopy software,” 5, Nat Methods 16(5), 387–395, Nature Publishing Group (2019) [doi:10.1038/s41592-019-0364-4].

15. T. Novák et al., “TestSTORM: Versatile simulator software for multimodal super-resolution localization fluorescence microscopy,” 1, Sci Rep 7(1), 951, Nature Publishing Group (2017) [doi:10.1038/s41598-017-01122-7].

16. M. Lindén et al., “Simulated single molecule microscopy with SMeagol,” Bioinformatics 32(15), 2394–2395 (2016) [doi:10.1093/bioinformatics/btw109].

17. V. Venkataramani et al., “SuReSim: simulating localization microscopy experiments from ground truth models,” 4, Nat Methods 13(4), 319–321, Nature Publishing Group (2016) [doi:10.1038/nmeth.3775].

18. M. Lagardère et al., “FluoSim: simulator of single molecule dynamics for fluorescence live-cell and super-resolution imaging of membrane proteins,” 1, Sci Rep 10(1), 19954, Nature Publishing Group (2020) [doi:10.1038/s41598-020-75814-y].

19. H. Heydarian et al., “Template-free 2D particle fusion in localization microscopy,” 10, Nat Methods 15(10), 781–784, Nature Publishing Group (2018) [doi:10.1038/s41592-018-0136-6].

20. D. Bourgeois, “Single molecule imaging simulations with advanced fluorophore photophysics,” 1, Commun Biol 6(1), 1–13, Nature Publishing Group (2023) [doi:10.1038/s42003-023-04432-x].

21. M. Hirsch et al., “A Stochastic Model for Electron Multiplication Charge-Coupled Devices – From Theory to Practice,” PLoS ONE 8(1), C.-T. Chen, Ed., e53671 (2013) [doi:10.1371/journal.pone.0053671].

22. Y. Zhang et al., “Investigating Single-Molecule Fluorescence Spectral Heterogeneity of Rhodamines Using High-Throughput Single-Molecule Spectroscopy,” J. Phys. Chem. Lett. 12(16), 3914–3921, American Chemical Society (2021) [doi:10.1021/acs.jpclett.1c00192].

23. B. Zhang, J. Zerubia, and J.-C. Olivo-Marin, “Gaussian approximations of fluorescence microscope point-spread function models,” Appl. Opt., AO 46(10), 1819–1829, Optica Publishing Group (2007) [doi:10.1364/AO.46.001819].

24. M. Ovesný et al., “ThunderSTORM: a comprehensive ImageJ plug-in for PALM and STORM data analysis and super-resolution imaging,” Bioinformatics 30(16), 2389–2390 (2014) [doi:10.1093/bioinformatics/btu202].

25. S. A. Mutch et al., “Deconvolving Single-Molecule Intensity Distributions for Quantitative Microscopy Measurements,” Biophysical Journal 92(8), 2926–2943, Elsevier (2007) [doi:10.1529/biophysj.106.101428].

26. T. J. Lambert, “FPbase: a community-editable fluorescent protein database,” 4, Nat Methods 16(4), 277–278, Nature Publishing Group (2019) [doi:10.1038/s41592-019-0352-8].

27. W. Sellmeier, “Ueber die durch die Aetherschwingungen erregten Mitschwingungen der Körpertheilchen und deren Rückwirkung auf die ersteren, besonders zur Erklärung der Dispersion und ihrer Anomalien,” Annalen der Physik 223(11), 386–403 (1872) [doi:10.1002/andp.18722231105].

28. B. P. Isaacoff et al., “SMALL-LABS: Measuring Single-Molecule Intensity and Position in Obscuring Backgrounds,” Biophysical Journal 116(6), 975–982 (2019) [doi:10.1016/j.bpj.2019.02.006].

29. Y. Zhang et al., “Minimizing Molecular Misidentification in Imaging Low-Abundance Protein Interactions Using Spectroscopic Single-Molecule Localization Microscopy,” Anal. Chem. 94(40), 13834–13841, American Chemical Society (2022) [doi:10.1021/acs.analchem.2c02417].

