## Supplementary material for "Physically informed Monte Carlo simulation of dual-wedge prism-based spectroscopic single-molecule localization microscopy": Fig. S

**Wei-Hong Yeo,[a] Cheng Sun,[b] Hao F. Zhang[a,*]**
[a] Northwestern University, Department of Biomedical Engineering, Evanston, United States
[b] Northwestern University, Department of Mechanical Engineering, Evanston, United States

***Hao F. Zhang,**

### Appendix A: Supplemental Material

**Table S1** Specifications and source of bandpass filters used for spectral calibration.

| Center wavelength (nm) | Full width at half maximum (nm) | Product Code | Manufacturer |
|---|---|---|---|
| 532 | 1 | FL532-1 | Thorlabs |
| 605 | 15 | FF01-605/15-25 | Semrock |
| 635 | 10 | FL635-10 | Thorlabs |
| 685 | 10 | 86-738 | Edmund Optics |
| 750 | 10 | FB750-10 | Thorlabs |

**Table S2** Optical and physical properties of Invitrogen microspheres used in the experiment.

| Excitation wavelength (nm) | Emission wavelength (nm) | Diameter of microspheres (μm) | Product Code |
|---|---|---|---|
| 540 | 560 | 0.1 | F8800 |
| 565 | 580 | 0.04 | F8794 |
| 580 | 605 | 0.1 | F8801 |
| 625 | 645 | 0.2 | F8806 |
| 633 | 720 | 0.04 | T8870 |

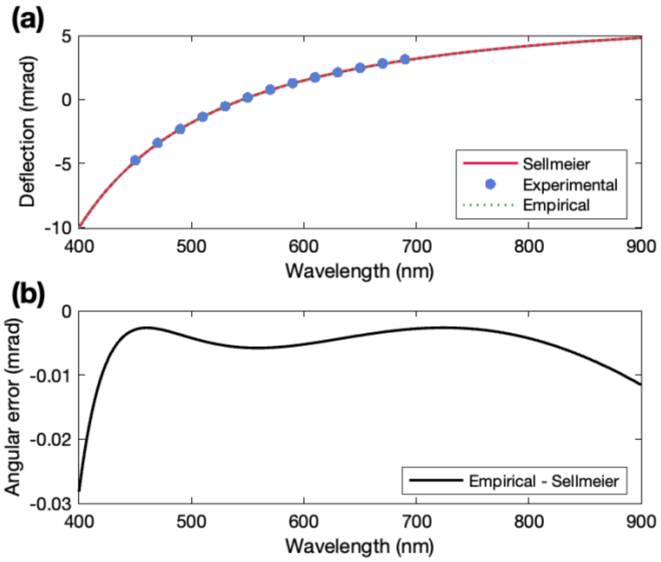


**Fig. S1** (a) Plot of the deflection of the DWP module determined experimentally (blue dots); theoretically using the Sellmeier equation (red solid line); and empirically using the rational best fit line (green dotted line). (b) Plot of the angular error between the empirical rational best fit line and the theoretical curve derived from Sellmeier equation.

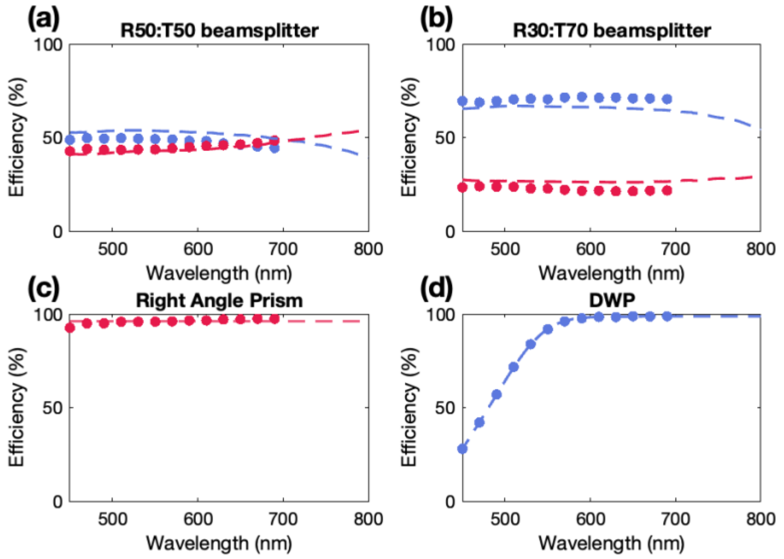


**Fig. S2** Plots of efficiency curves of (a) R50:T50 beamsplitter, (b) R30:T70 beamsplitter, (c) right-angle prism, (d) dual wedge prism. We obtained efficiency data using a supercontinuum laser with an AOTF setup and shown our data as dots in the graph. The dotted lines for (a)-(b) refer to the Thorlabs datasheet; for (c), the average over the experimental datapoints acquired; and for (d), the extrapolated curve from the experimental datapoints acquired. In all graphs, blue datapoints refer to transmission efficiencies, and red datapoints refer to reflection efficiencies.

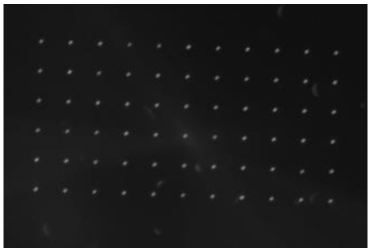

**Fig. S3** A sample image of our nanohole array.
